# Exploring the kinetic selectivity of antipsychotics for dopamine D_2_ and 5-HT_2A_ receptors: implications for the prevalence of EPS and receptor occupancy

**DOI:** 10.1101/2021.11.14.468520

**Authors:** David A. Sykes, Jack Lochray, Hannah M.F. Comfort, Palash Jain, Steven J. Charlton

## Abstract

Certain atypical antipsychotic drugs (APDs) used in the treatment of schizophrenia have been hypothesized to show reduced extrapyramidal side effects (EPS), due to their ability to promote nigrostriatal dopamine release through 5-HT_2A_ receptor (5-HT_2A_R) blockade. The strength of this hypothesis is currently limited to a consideration of the relative receptor affinities of APDs for the 5-HT_2A_R and dopamine D_2_ receptor (D_2_R). Here we measure the 5-HT_2A_R kinetic binding properties of a series of typical and atypical APDs in a novel time-resolved fluorescence resonance energy transfer assay and correlate these properties with their observed EPS at therapeutic doses. For compounds with negligible affinity for 5-HT_2A_R, EPS is robustly predicted by a D_2_R specific rebinding model that integrates D_2_R association and dissociation rates to calculate the net rate of reversal of receptor blockade (*k*_r_). However, we show that for compounds with significant affinity for the 5-HT_2A_R, such as sertindole, higher relative 5-HT_2A_ occupancy over time is an indicator for a reduced propensity to cause EPS.

This study suggests that there is room for the development of novel kinetically optimised antipsychotic agents that modulate both serotonergic and dopamine function in a manner beneficial in the treatment of this chronic and debilitating disease.

## Introduction

The common mechanism of antipsychotic drug action is postsynaptic dopamine antagonism (Creese et al., 1975; Seeman and Lee, 1975; Seeman et al., 1976), which forms the basis of the original ‘dopamine hypothesis’ of schizophrenia and the subsequent ‘modified dopamine hypothesis’ which argued for the regional specificity of dopamine signalling and drug action (Carlsson and Lindqvist, 1963; Davis et al., 1991). However other signalling pathways have been suggested to play a key role in both the therapeutic and favourable side effect profile of second-generation atypical antipsychotic drugs (APDs) over conventional, first-generation, or typical APD treatments (Karam et al., 2010; Meltzer, 1999). Atypical APDs are those that produce a lower incidence of side effects at clinically effective doses (Meltzer, 1999).

In truth, APDs rarely fall neatly into one of these two classes since their side effects comprise multiple, pharmacologically separable events, reflective of their pharmacological heterogeneity (Kusumi et al., 2015; Mackin and Thomas, 2011). Consequently, a more rational approach to APD classification and drug design is to consider a continuum (or scale) based on the specific pharmacology of a given side effect, rather than the current dichotomous system. Second-generation atypical APDs can be assigned to three distinct subclasses based on affinity; those considered relatively specific for dopamine receptors, (the so-called dopamine D_2_/D_3_ APDs e.g. amisulpride, sulpiride and remoxipride), serotonin-dopamine antagonists (e.g. risperidone, sertindole and iloperidone), and finally multi-acting receptor-targeted antipsychotics or MARTAS (e.g. clozapine, olanzapine, asenapine, ziprasidone and quetiapine) which share properties of the second class but in addition may display direct effects at other receptors (e.g. 5-HT_1A_ agonism). A new third generation of antipsychotics (e.g., aripiprazole) has emerged characterised by their high dopamine D_2_ affinity, low intrinsic activity or partial D_2_ agonism (Mailman and Murthy, 2010).

Meltzer and colleagues were first to label simultaneous 5-HT_2A_ and D_2_ receptor blockade as a pharmacodynamic (PD) mechanism which differentiates out the clinical effect of typical and atypical agents (Meltzer et al., 1989). In their seminal paper they defined atypicality using two criteria: firstly, as drugs with greater affinity for the 5-HT_2A_ receptor over D_2_ receptor and secondly as a reduced affinity for D_2_Rs than seen for conventional APs.

Serotonin neurones innervate dopamine neurones either directly via postsynaptic 5-HT_2A_Rs on the dopamine neurone, or indirectly on GABA interneurons (Celada et al., 2013; de Almeida and Mengod, 2007). 5-HT has been proposed to act at these 5-HT_2A_Rs and inhibit the release of dopamine whereas antagonists have been proposed to increase release (Horacek et al., 2006; Lucas and Spampinato, 2000; Stahl, 2008; Stahl, 2013). The opposite occurs at 5-HT_1A_ autoreceptors present on 5-HT neurones, with activation thought to lead to a reduction in 5-HT release and a subsequent reduction in 5-HT_2A_ postsynaptic receptor activation and as a consequence accelerated (or disinhibited) dopamine release (Horacek et al., 2006; Stahl, 2008; Stahl, 2013).

It has been suggested that tissue-selective antagonism of D_2_/5-HT_2A_ receptors by atypical APDs may play a role in both their improved efficacy against negative symptoms but also their reduced side effect profile (Leucht et al., 2013; Leucht et al., 2009; Meltzer, 1992; Meltzer et al., 1989; Schotte et al., 1996; Sulpizio et al., 1978). In terms of the on-target side effect profile of clinically used APDs a model has been proposed whereby antagonism of 5-HT_2A_Rs in the cortex results in increased dopamine release from neurons in the striatum reducing striatal APD occupancy and decreasing the subsequent risk of EPS (reviewed in (Kapur and Remington, 1996) and (Schotte et al., 1996). What is missing from this purely affinity-based model is a consideration of the kinetic properties of the APDs at both the serotonergic 5-HT_2A_ and dopamine D_2_Rs and the influence this could have on D_2_R occupancy over time.

We recently established an alternative mechanism of atypicality based on a restricted drug diffusion model, which suggests that for the majority of compounds association rate and not dissociation rate better predicts the overall rate of dopamine D_2_ receptor reversal in the striatum through a process of receptor rebinding (Sykes et al., 2017). In the aforementioned study the correlation of dopamine D_2_ receptor binding reversal rate (*k*_r_) with clinically derived EPS was not perfect and there were certain compounds whose EPS profile was not so well described by this model leading us to speculate that the ‘serotonin-dopamine hypothesis’ at least in terms of EPS profile may have some validity for compounds with significant affinity for 5-HT_2A_Rs.

The overall aim of this study is to consider the role of 5-HT_2A_Rs and receptor binding kinetics in dictating the side effect profile of clinically used antipsychotic agents and in doing so explore the relevance to the ‘serotonin-dopamine hypothesis’ and possible strategies for the development of novel kinetically optimised antipsychotic agents that modulate both serotonergic and dopamine function in a manner beneficial to the treatment of this chronic and debilitating disease.

## Methods

### Materials

Tag-lite labeling medium (LABMED), SNAP-Lumi4-Tb and the CellAura fluorescent 5-HT_2A/2B_ receptor antagonist [4F4PP oxalate-red] (CA201019), was obtained from Hello Bio (Bristol, UK). 96-well polypropylene plates (Corning) were purchased from Fisher Scientific UK (Loughborough, UK) and 384-well Optiplate plates were purchased from Perkin Elmer (Beaconsfield, UK). 5’-Guanylyl imidodiphosphate (GppNHp), risperidone, chlorpromazine hydrochloride, quetiapine hemifumarate, ziprasidone hydrochloride monohydrate, ketanserin (+)-tartrate, zotepine, sertindole and spiperone used in competition assays were obtained from Sigma-Aldrich (Poole, UK). Olanzapine, clozapine, paliperidone, amisulpride, asenapine maleate and haloperidol hydrochloride used for competition assays were obtained from Tocris Bioscience (Avonmouth, Bristol). Lurasidone and Iloperidone were obtained from Selleckchem.

### Cell culture

The host Chinese hamster ovary (CHO) K1 cell line was transiently transfected using Fugene (Fugene: DNA ratio 3:1) with the cDNA encoding a SNAP-tagged human serotonin 5-HT_2A_ receptor (Cis Bio, France). CHO-5HT_2A_ cells were maintained in Dulbecco’s modified Eagle’s medium: Ham F12 (DMEM: F12) containing 2mM glutamine (Sigma-Aldrich, Poole, UK) and supplemented with 10% fetal calf serum (Life Technologies, Paisley UK). CHO-D_2L_ transfected with the cDNA encoding a SNAP-tagged human receptors from Genbank (ref: NM_000795) and CisBio respectively, were grown as previously described (Sykes et al., 2017).

### Terbium labeling of SNAP-tagged cells

Cell culture medium was removed from the t175cm^2^ flasks containing confluent adherent CHO-5HT_2A_ 12 mL of Tag-lite labeling medium containing 100 nM of SNAP-Lumi4-Tb was added to the flask and incubated for 1 h at 37 °C under 5% CO_2_. Cells were washed 2x in PBS (GIBCO Carlsbad, CA) to remove the excess of SNAP-Lumi4-Tb then detached using 5 mL of GIBCO enzyme-free Hank’s-based cell dissociation buffer (GIBCO, Carlsbad, CA) and collected in a vial containing 5mL of DMEM: F12 containing 2mM glutamine (Sigma-Aldrich) and supplemented with 10% fetal calf serum. Cells were pelleted by centrifugation (5 min at 1500 rpm) and the pellets were frozen to −80 °C. To prepare membranes homogenisation steps were conducted at 4°C (to avoid receptor degradation). Specifically, 20 ml per t175-cm^2^ flask of wash buffer (10 mM HEPES and 10 mM EDTA, pH 7.4) was added to the pellet. This was homogenized using an electrical homogenizer Ultra-Turrax (Ika-Werk GmbH & Co. KG, Staufen, Germany) (position 6, 4 × 5-s bursts) and subsequently centrifuged at 48,000*g* at 4°C (Beckman Avanti J-251 Ultracentrifuge; Beckman Coulter, Fullerton, CA) for 30 min. The supernatant was discarded, and the pellet was re-homogenized and centrifuged as described above in wash buffer. The final pellet was suspended in ice-cold 10 mM HEPES and 0.1 mM EDTA, pH 7.4, at a concentration of 5 to 10 mg/ml. Protein concentration was determined using the bicinchoninic acid assay kit (Sigma-Aldrich), using BSA as a standard and aliquots maintained at −80°C until required. Prior to their use, the frozen membranes were thawed, and the membranes suspended in the assay buffer at a membrane concentration of 0.2mg/ml.

### Fluorescent ligand-binding assays

5-HT_2A_ fluorescent binding experiments employed 4F4PP oxalate-red respectively whilst PPHT-red was employed in the case of the human dopamine D_2_R. Experiments were conducted in white 384-well Optiplates, in assay binding buffer, HBSS containing 5 mM HEPES, 0.02% pluronic acid, 1% DMSO pH 7.4 and 100 _μ_M GppNHp. GppNHp was included to remove the G protein–coupled population of receptors that can result in two distinct populations of binding sites in membrane preparations, since the Motulsky–Mahan model (1984) is only appropriate for ligands competing at a single site. In the case of the human 5-HT_2A_R, nonspecific binding was determined in the presence of sertindole (10 µM), and D_2L_ specific PPHT-red ligand binding was performed in the presence of haloperidol (10 µM).

### Determination of tracer binding kinetics

To accurately determine 4F4PP oxalate-red association rate (*k*_on_) and dissociation rate (*k*_off_) values, the observed rate of association (*k*_ob_) was calculated using at least six different concentrations. The appropriate concentration of 4F4PP oxalate-red was incubated with human 5-HT_2A_ CHO cell membranes (4 µg per well) in assay binding buffer (final assay volume, 40µl). The degree of 4F4PP oxalate-red bound to the receptor was assessed at multiple time points by HTRF detection to allow construction of association kinetic curves. The resulting data were fitted to a one phase exponential association model, equation (2) to derive an estimate of *k*_on_ and *k*_off_ from plotting tracer concentration versus observed association (*k*_obs_) as described under Data Analysis. F-PPHT-red tracer binding was carried out as previously described (see Sykes et al. (2017)).

### Competition binding kinetics

To determine the association and dissociation rates of 5-HT_2A_R specific ligands, we used a competition kinetic binding assay. This approach involves the simultaneous addition of both fluorescent ligand and competitor to the receptor preparation, so that at t = 0 all receptors are unoccupied. 50 nM 4F4PP oxalate (a concentration which avoids ligand depletion in this assay volume, see (Carter et al., 2007) was added simultaneously with the unlabeled compound (at t = 0) to CHO cell membranes containing the human 5-HT_2A_R (4 µg per well) in 40 µl of assay buffer. The degree of 4F4PP oxalate-red bound to the receptor was assessed at multiple time points by HTRF detection. Similarly, D_2_R specific ligands were profiled as previously described (see Sykes et al. (2017)).

Nonspecific binding was determined as the amount of HTRF signal detected in the presence of sertindole (10 µM) in the case of the 5-HT_2A_R and was subtracted from each time point, meaning that t = 0 was always equal to zero. Each time point was conducted on the same 384 well plate incubated at 37°C with orbital mixing (1sec of 100 RPM/cycle). Multiple concentrations of unlabeled competitor were tested, for determination of rate parameters. Data were globally fitted using equation (3) to simultaneously calculate *k*_on_ and *k*_off_. The estimation of lurasidone and iloperidone binding kinetics at the dopamine D_2_R was carried out as previously described (Sykes et al., 2017).

### Signal detection and data analysis

Signal detection was performed on a Pherastar FS (BMG Labtech, Offenburg, Germany) using standard HTRF settings. The terbium donor was always excited with 3 laser flashes at a wavelength of 337 nm. A kinetic TR-FRET signal was collected at 20 seconds intervals both at 665 nm and 620 nm, when using red acceptor. HTRF ratios were obtained by dividing the acceptor signal (665 nm) by the donor signal (620 nm) and multiplying this value by 10,000. Probe dissociation rates were analysed by displacement of the tracer with a large excess of an unlabelled ligand known to bind to the same site with similar or higher affinity.

All experiments were analyzed by non-regression using Prism 6.0 (GraphPad Software, San Diego, U.S.A.). Competition displacement binding data were fitted to sigmoidal (variable slope) curves using a “four parameter logistic equation”:

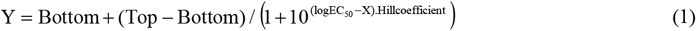

IC_50_ values obtained from the inhibition curves were converted to *K*_i_ values using the method of (Cheng and Prusoff, 1973). 4F4PP oxalate-red association data was analysed using nonlinear regression to fit the specific binding data to a one phase exponential association equation.

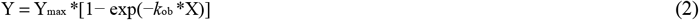

Where X is time (min) and *k*_ob_ is the observed rate of association (min^-1^). 4F4PP oxalate-red binding followed a simple law of mass action model, with *k*_ob_ increasing in a linear fashion as a function of tracer concentration up to a value of 50nM. The slope of the line equates to the association rate (*k*_on_), and extrapolation of the plot to y = 0 yields the dissociation rate (*k*_off_) (Sykes et al., 2010).

Association and dissociation rates for unlabeled antagonists were calculated using the equations described by (Motulsky and Mahan, 1984):

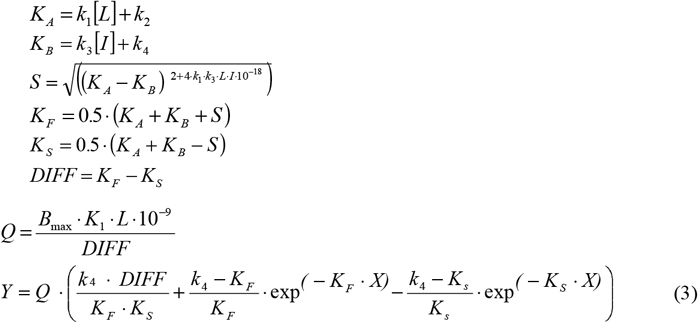

Where: X = Time (min), Y = Specific binding (HTRF ratio 665nm/620nm*10,000), *k*_1_ = k_on_ tracer-red, *k*_2_ = k_off_ tracer-red, L = Concentration of 4F4PP oxalate-red used (nM), B_max_ = Total binding (HTRF ratio 665nm/620nm*10,000), I = Concentration of unlabeled antagonist (nM). Fixing the above parameters allowed the following to be calculated: k_3_ = Association rate of unlabeled ligand (M^-1^ min^-1^), k_4_ = Dissociation rate of unlabeled ligand (min^-1^). Dissociation experiments were fitted to a one phase mono-exponential decay function to estimate the dissociation rate of 4F4PP oxalate-red directly. Specific binding was determined by subtracting the nonspecific HTRF ratio from the total HTRF ratio.

### 5-HT_2A_ calcium assay

The ability of the antipsychotics sertindole and clozapine to block 5-HT_2A_ receptor-linked G-protein activation was assessed through 5-HT stimulated Ca2+ release. Briefly, CHO-h5-HT_2A_ cells (40,000 per well) were plated into black 96-well plates and grown overnight in F-12: DMEM containing 10% dialysed FBS. Intracellular calcium was detected using a FLIPR calcium 5 no wash assay kit and read on a FlexStation 3 plate reader (Molecular Devices, Sunnyvale USA).

Cells were incubated in duplicate with the antipsychotics sertindole and clozapine at 10, 30 and 100 x*K*_d_ based on their respective binding affinities at the 5-HT2AR for 60min. Dye and compounds were made up in assay buffer (HBSS containing 20mM HEPES and 0.1%(w/v) BSA (pH 7.4)) and added directly to cells in media.

Activation of CHO-h5-HT_2A_Rs was initiated through application of the endogenous agonist 5-HT and calcium responses were monitored for a period of 90seconds. Maximal calcium responses were determined by calculating maximal response – minimal response (Max-Min).

### Modelling drug-receptor re-binding at Dopamine D_2_ and serotonin 5-HT_2A_ receptors

Rebinding describes the ability of a drug molecule to bind to multiple receptors within a compartment before diffusing away into bulk, the overall effect being extended target-receptor occupancy (Vauquelin and Charlton, 2010). To examine this we utilized a model of an immunological synapse with a compartment volume of 0.176 _μ_m^3^, which is within the range described for the dopamine synapse (Coombs and Goldstein, 2004). In this model, the overall macroscopic reversal rate (*k*_r_) is described by the following equation;

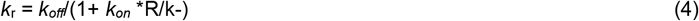

where: *k*_off_ = dissociation rate from the receptor, *k*_on_ = association rate onto the receptor, [R] = surface receptor density fixed at 1×10^11^ cm^-2^ for D_2_R calculations and 3×10^11^ cm^-2^ for 5-HT_2A_R calculations, and k-= the diffusion rate out of the synaptic compartment into bulk aqueous, fixed at 1.2×10^-5^ cm/sec.

Based on this rebinding model the effectiveness of APDs to reverse dopamine D_2_ receptor blockade and therefore relieve the symptoms of EPS via their effects at 5-HT_2A_Rs was estimated. This was achieved by calculating the relative affinities of clinically used APDs for 5-HT_2A_: D_2_ receptors, and through determining area under the curve (AUC) estimates of 5-HT2A and D_2_R relative receptor occupancy (5-HT_2A_: D_2_) at D_2_R occupancy levels between 80-65% (i.e. those relevant to antipsychotic and EPS through combined serotonergic and D_2_R blockade see (Kasper et al., 1999). AUC estimates of 5-HT_2A_ and D_2_R relative receptor occupancy (5-HT_2A_: D_2_) were assessed utilising previously reported human D_2L_R specific kinetic values (Sykes et al., 2017) and 5-HT_2A_ values reported herein.

In all cases initial dopamine D_2_R striatal occupancy was assumed to reach 80% with individual 5-HT_2A_R starting occupancies varied based on the relative affinities of these clinically used antipsychotics for the D_2_R and 5-HT_2A_R. The resulting AUC values are an integrated measurement of receptor occupancy based on a rebinding phenomenon and are used as a cumulative measurement of drug effect. All data were analyzed using GraphPad Prism 6.0.

### Comparing binding kinetics and clinical side effect profile

To explore the role of kinetics in determining on-target side effect liability we correlated the kinetic values determined in this study with published clinical data taken from a comprehensive meta-analysis of clinically used APDs performed by Leucht et al., (2013). Unless otherwise stated correlation analyses were performed using a two-tailed Spearman rank correlation allowing the calculation of the correlation coefficient, r_s_. Although this analysis does not assume a linear relationship, a simple trend line has been added to illustrate the positive or negative association between the two variables. Differences were considered significant at P< 0.05. All analysis were performed using GraphPad Prism 6.0.

## Results

### Characterization of 4F4PP oxalate-red binding to the human 5-HT_2A_ receptor

Specific binding of the antagonist 4F4PP oxalate-red to human 5-HT_2A_Rs expressed in CHO membranes was saturable and best described by the interaction of the fluorescent ligand with a single population of binding sites (Figure. 1A). Non-specific binding was linear and represented just over 50% of total binding. This could reflect the relatively high log D of the parent molecule (3.63) at physiological pH 7.4, which suggests it may penetrate biological membranes creating the potential for increased non-specific proximity. From these studies, the equilibrium dissociation constant (*K*_d_) of 4F4PP oxalate-red was determined to be 3.98 ± 0.65 nM, n=23.

**Figure 1.**
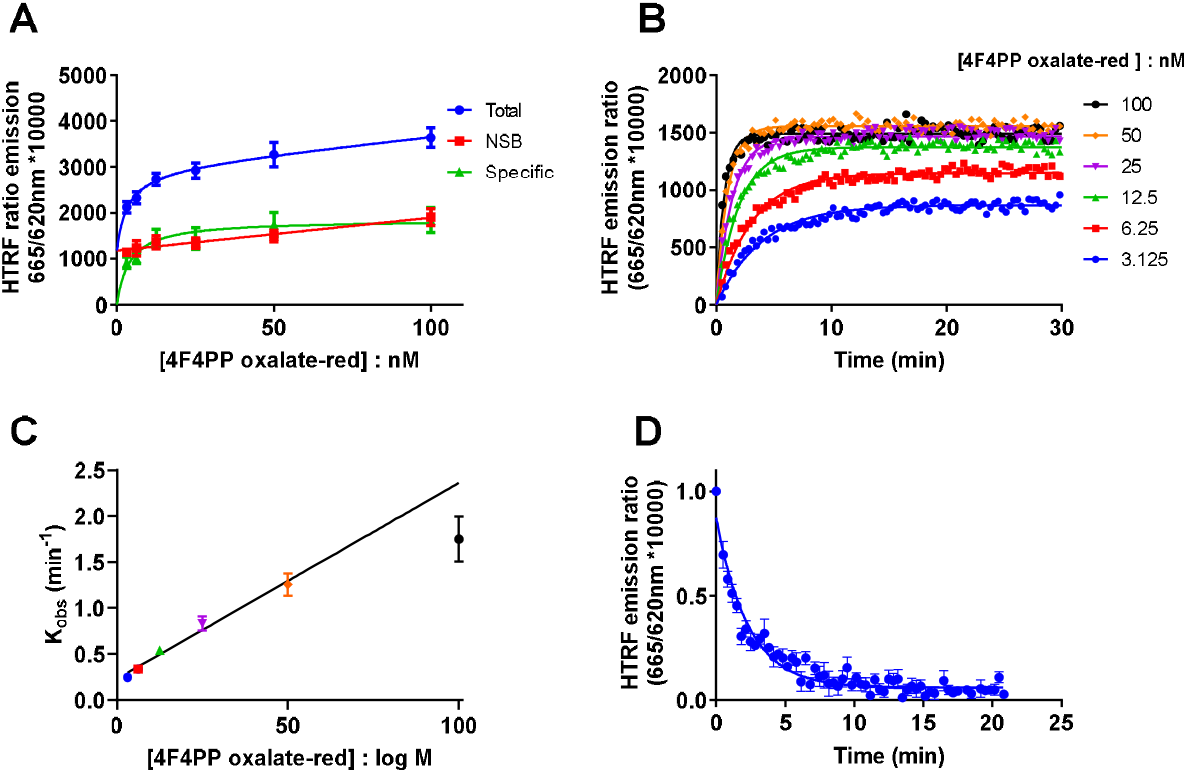
Determination of 4F4PP oxalate-red equilibrium and kinetic binding parameters. (**A**) Saturation analysis showing the binding of 4F4PP oxalate-red to the human 5-HT_2A_ receptor. CHO-5-HT_2A_ receptor cell membranes (4 µg per well) were incubated for 60 min with gentle agitation with increasing concentrations of 4F4PP oxalate-red. Data are presented in duplicate form from a representative of 23 separate experiments. (**B**) The observed association of 4F4PP oxalate-red to the human 5-HT2A receptor. Data are presented as the mean from 24 separate experiments. (**C**) A plot of 4F4PP oxalate-red concentration versus *k*_obs_. Data are presented as mean ± SEM from a total of 24 separate experiments. *k*_obs_ data for the highest concentration of fluorescent ligand tested (100nM, >30x *K*_d_), is shown but was excluded from the analysis. (**D**) 4F4PP oxalate-red dissociation following the addition of sertindole (10 µM). Dissociation data are presented in mean ± SEM from 4 experiments performed in singlet. All binding reactions were performed in the presence of GppNHp (0.1mM) with non-specific binding levels determined by the inclusion of sertindole (10 µM).

The binding kinetics of 4F4PP oxalate-red was characterised by monitoring the observed association rates at six different ligand concentrations (Figure 1B). Its observed rate of association was related to 4F4PP oxalate-red concentration in a linear fashion up to a concentration of 50nM (>10x *K*_d_). The highest concentration of fluorescent ligand employed (100nM, >30x *K*_d_), on average showed a variable and inconsistent deviation in *k*_obs_ creating an apparent non-linear relationship.

4F4PP oxalate-red kinetic rate parameters for 4F4PP oxalate-red were calculated based on a limited concentration range (3.12-50 nM) from a global fit of association curves to derive a single best-fit estimate for *k*_on_ and *k*_off_, resulting in a *k*_on_ estimate of 4.21 ± 0.47 x10^7^ M^-1^ min^-1^ and a *k*_off_ of 0.09 ± 0.01 min^-1^, n= 24, see Figure 1B. The resulting *K*_d_ (*k*_off_ / *k*_on_) of 2.82 ± 0.47 nM was comparable to that obtained from the equilibrium studies described above.

In addition, the kinetic rate parameters for 4F4PP oxalate-red (3.12-50 nM) were calculated from plots of tracer concentration versus observed rates of association (*k*_obs_), resulting in a *k*_on_ estimate of 2.37 ± 0.27 x10^7^ M^-1^ min^-1^ and a *k*_off_ of 0.25 ± 0.03 min^-1^, n= 24, see Figure 1C. The resulting *K*_d_ (*k*_off_ / *k*_on_) of 13.1 ± 1.6 nM was significantly different from that obtained from the equilibrium saturation binding studies described above (Bonferroni’s multiple comparisons test, P<0.0001). Ligand dissociation estimated directly through the addition of an excess of sertindole revealed a *k*_off_ value of 0.36 ± 0.06 min^-1^, see Figure 1D.

### Characterization of unlabeled APDs binding to the human 5-HT_2A_ receptor

The binding affinity of the various ligands for the human 5-HT_2A_R was measured at equilibrium at 37°C in a buffer containing GppNHp (0.1 mM) to ensure that antagonist binding only occurred to the G protein-uncoupled form of the receptor. 5-HT2AR binding affinities (*K*_i_ values) for the APDs studied are summarized in Table 1, and competition curves are presented in Figure. 2A and B. In Table 1 APDs are separated into different groups based on their recognised pharmacology and those described in the literature as typical or atypical and those described as both. Dopamine D_2L_ receptor-specific kinetics parameters for lurasidone and iloperidone are summarised in Supplemental Table 1.

**Table 1.**
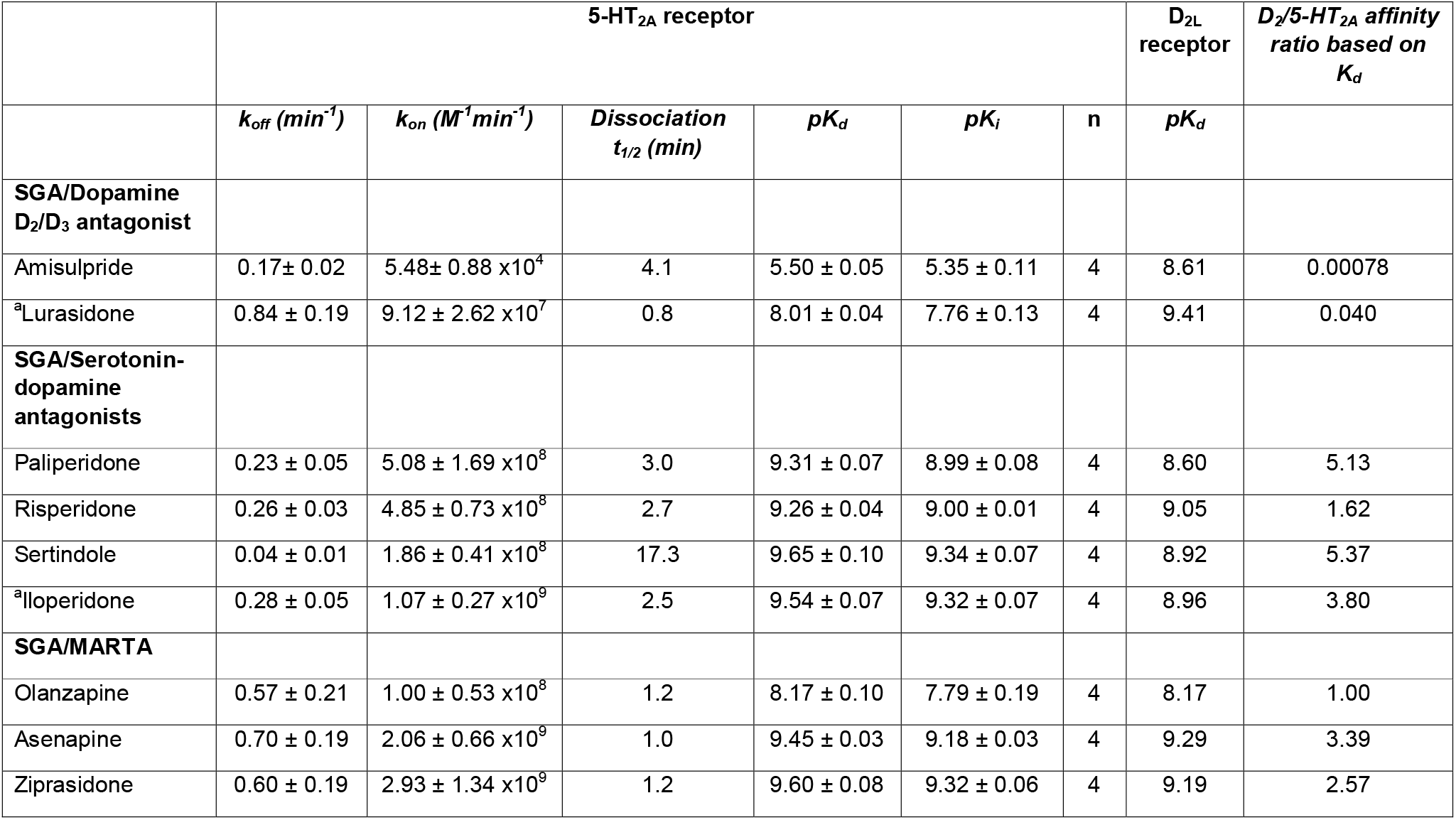
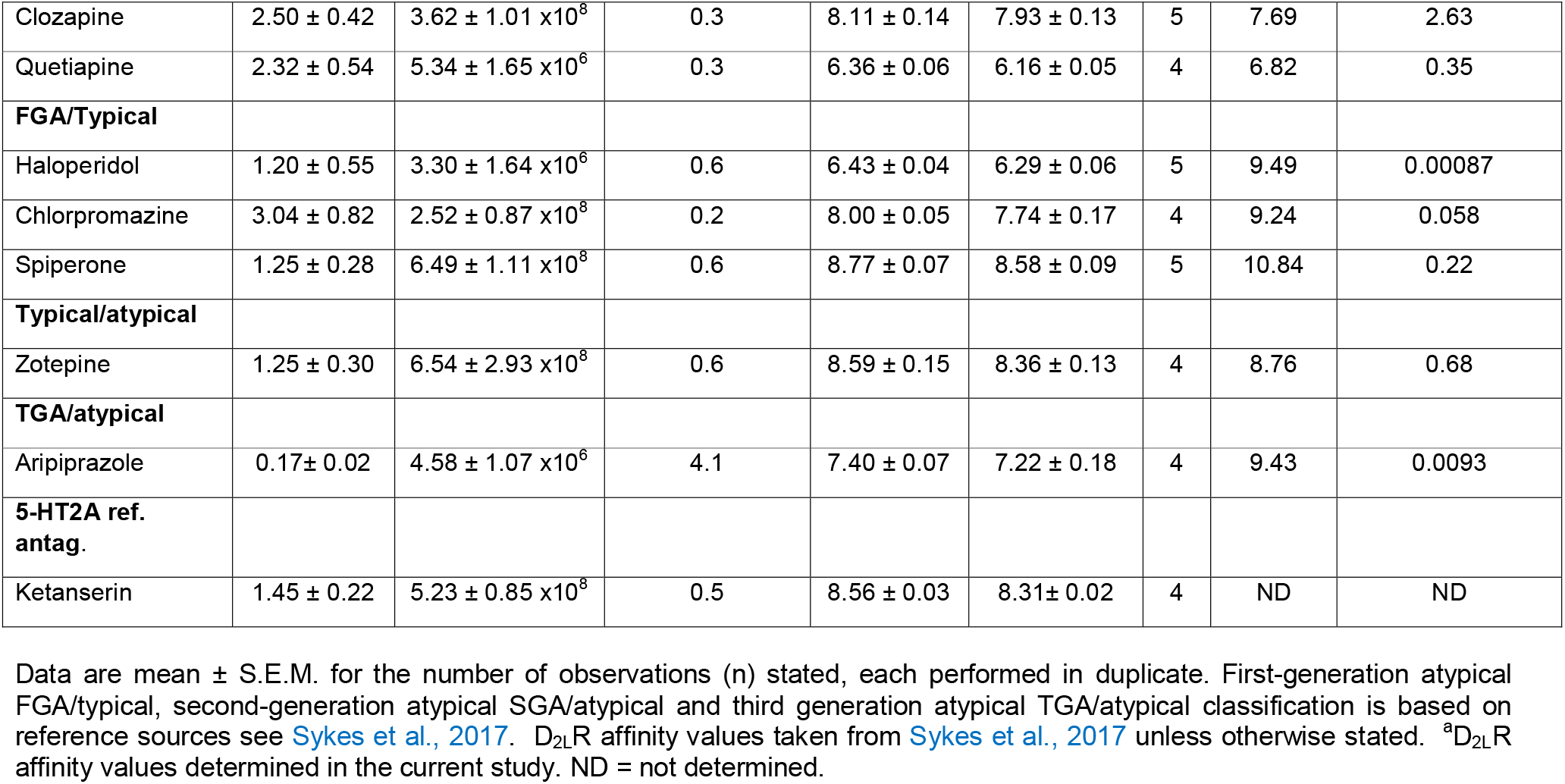
Kinetic binding parameters of unlabelled clinically used APDs and reference antagonist for human 5-HT_2A_ receptors and their historical classification as atypical or typical and those characterized as typical/atypical.

**Figure 2.**
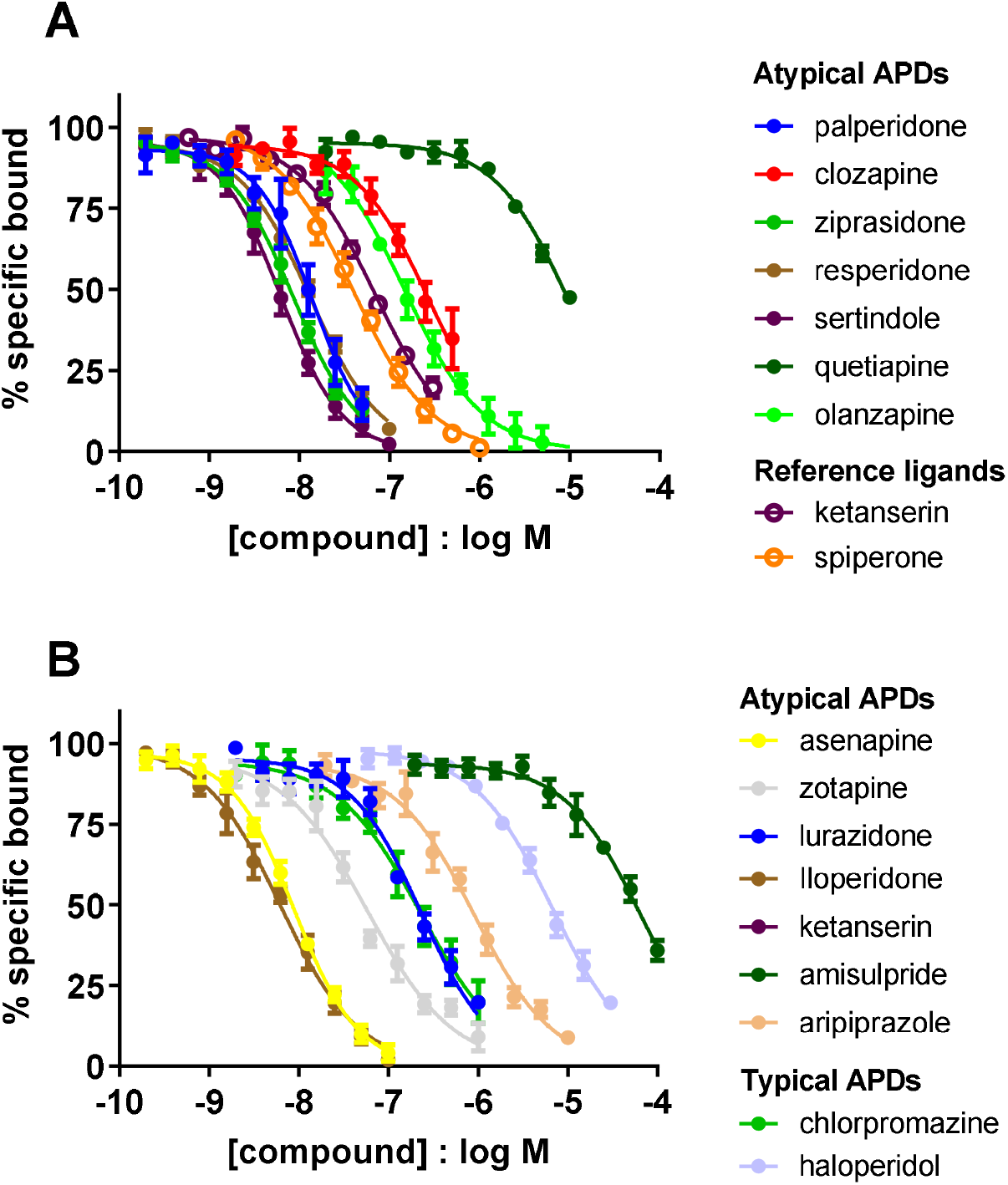
Equilibrium competition binding for APDs and reference ligands at the human 5-HT _2A_ receptor. **(A)** and **(B)** Competition between 4F4PP oxalate-red (50 nM) and increasing concentrations of atypical APDs, typical APDs and 5-HT_2A_ reference ligands for the human 5-HT_2A_ receptor. All binding reactions were performed in the presence of GppNHp (0.1mM) with non-specific-binding levels determined by the inclusion of sertindole (10 _μ_M). Equilibrium data were fitted to the equations described in the methods to calculate *K*_i_, values for the unlabeled ligands; these are summarized in Table 1. Data are presented as mean ± S.E.M. values from four or five separate experiments performed in duplicate. The number of observations (n) for each ligand is stated in Table 1. All data used in these plots are detailed in Table 1. *Zotepine was originally introduced as an atypical APD but is sometimes described as a typical APD, based on its side effects profile. Classification is based on reference sources see Sykes et al., 2017.

Representative 4F4PP oxalate-red competition association binding curves for selected D_2_R ligands are shown in Figure 3A-E. Progression curves for 4F4PP oxalate-red alone and in the presence of competitor were fitted to equation 3, enabling the calculation of both *k*_on_ (k3) and *k*_off_ (k4) for each of the ligands, as reported in Table 1.

**Figure 3.**
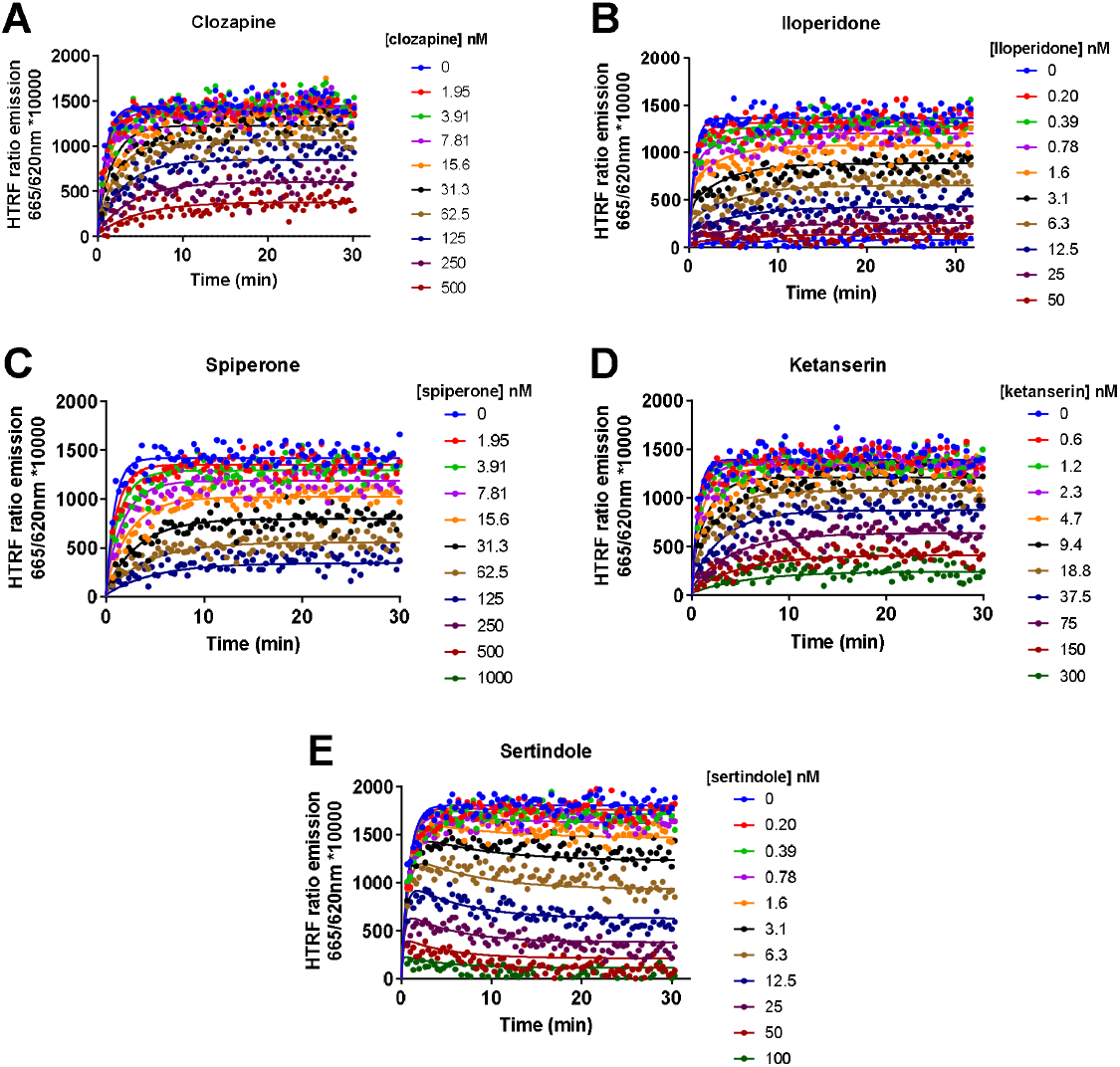
Competition association binding at the 5-HT_2A_ receptor. Competition between 4F4PP oxalate-red (50 nM) and increasing concentrations of representative atypical and typical APDs **(A)** clozapine, **(B)** iloperidone, **(C)** spiperone, **(D)** ketanserin and **(E)** sertindole at the human 5-HT_2A_ receptor. All binding reactions were performed in the presence of GppNHp (0.1mM) with non-specific-binding levels determined by the inclusion of sertindole (10 _μ_M). Kinetic and equilibrium data were fitted to the equations described in the methods to calculate *K*_i_, *K*_d_, and *k*_on_ and *k*_off_ values for the unlabeled ligands; these are summarized in Table 1. Data are presented as mean ± S.E.M. values from at least 3 or more separate experiments performed in duplicate. The number of observations (n) for each ligand is stated in Table 1. All data used in these plots are detailed in Table 1.

There was a very wide range of dissociation rates for the different ligands, with dissociation half-lives (t_½_) (where, t_½_ = 0.693/ *k*_off_) between 0.2 min for chlorpromazine to 17.3 min for sertindole. Confidence intervals on the kinetic fits for sertindole were on average much smaller, compared to the rapidly dissociating chlorpromazine, reflective of both the tracers own kinetic properties and the off-line addition protocol chosen for these studies. To validate the rate constants, the kinetically derived dissociation constant (*K*_d_) values (*k*_off_ / *k*_on_) were compared with the dissociation constant (*K*_i_) obtained from equilibrium competition binding experiments (see Supplementary Figure 1). There was a very good correlation between these two values for all APDs tested (Two-tailed Pearson’s correlation r^2^ = 0.99, P <0.0001) indicating the kinetic parameters were consistent with the equilibrium derived affinity values. Previous radioligand binding studies have reported dissociation rates for [^3^H]-ketanserin in the rat of 0.7 min^-1^ (t_1/2_ = 1min) in a salt containing buffer at 37°C which is similar (2-fold slower) to the values we report in our current study (Leysen et al., 1982). Any differences observed between this and the original studies could potentially have arisen from the use of different species (rat cortical tissue), or the use of a more alkaline buffer of lower salt concentration (eg. 50mM Tris-HCl, 120mM NaCl pH7.7) (Leysen et al., 1982; Schotte et al., 1983).

### Functional consequences of slow dissociation from 5-HT2A receptors

Like our previous study comparing the binding of APDs to the hD_2L_R we observed a wide range in *k*_on_ values between the APDs (>10,000-fold) and a relatively wide range in *k*_off_ values (∼70-fold) for ligands binding to the human 5-HT_2A_R. In contrast to our previous dopamine D_2L_R study (Sykes et al.,2017), the typical APDs examined here exhibited a much wider variation in the value of *k*_on_, driving differences in 5-HT_2A_R affinity. More variation in both on and off-rates were observed for atypical APDs, notably, sertindole exhibited a *k*_off_ value much slower than any other clinically used atypical APDs tested (Table 1).

Calcium assays are useful bioassays to assess the kinetics of antagonist binding since slowly dissociating antagonist readily suppress calcium signalling leading to insurmountable reductions in measurable agonist response. This occurs since calcium assays are by nature non-equilibrium assays with agonist responses occurring over a short time frame (seconds) and usually insufficient for the binding of agonist and antagonist to reach a true equilibrium (Charlton and Vauquelin 2010; Bdioui et al., 2018). The ability of sertindole to display insurmountable antagonism at 5-HT_2A_Rs in response to increasing concentrations of 5-HT in a calcium release assay at physiological temperature is highlighted in Figure 4B. In contrast, clozapine’s effects are almost fully reversible reflecting its relatively rapid off-rate from 5-HT_2A_Rs Figure 4A.

**Figure 4.**
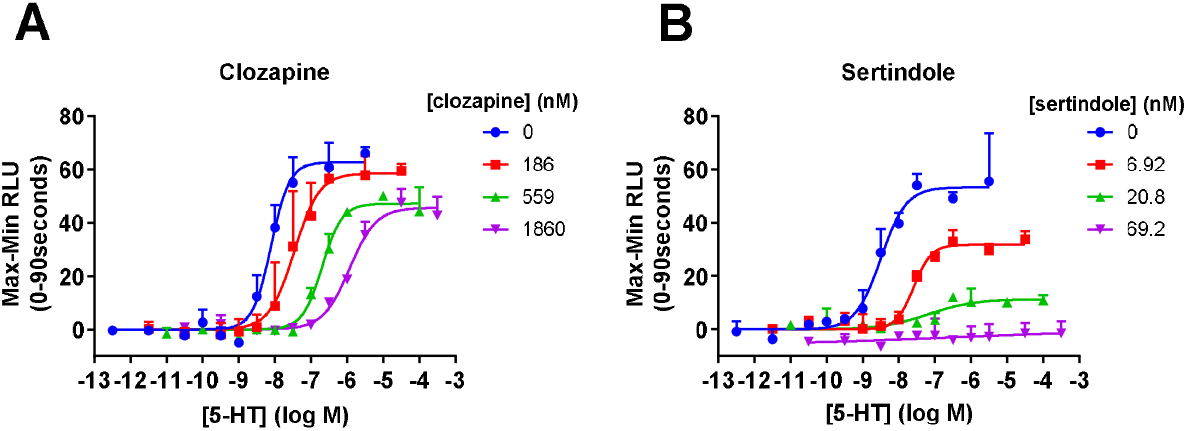
5-HT concentration calcium responses curves obtained in CHO-5-HT_2A_ cells in the absence and presence of the atypical APDs **(A)** clozapine and **(B)** sertindole at 37°C. Calcium data are presented as mean ± range from a representative of four separate experiments performed in duplicate.

### Comparing APD 5-HT2A and dopamine D_2_ receptor affinity

In consideration of a role for 5-HT_2A_Rs in the reduced incidence of EPS it has been widely reported that the 5-HT_2A_ affinity of certain clinically used atypical antipsychotics is much higher than their corresponding affinities at dopamine D_2_Rs (Horacek et al., 2006; Kusumi et al., 2015; Meltzer et al., 1989; Richelson and Souder, 2000). For example, estimates for clozapine taken directly from these papers range from 14 to 80-fold higher affinity for the 5-HT_2A_R over the D_2_ receptor. A comparison of the equilibrium and kinetic parameters of these compounds for these two receptors is shown in Figure 5. The TR-FRET data revealed a surprisingly small difference in affinity for D_2_ and 5-HT_2A_Rs across the clinically used atypical serotonin-dopamine antagonists (e.g. risperidone, sertindole and iloperidone) and MARTAs studied.

**Figure 5.**
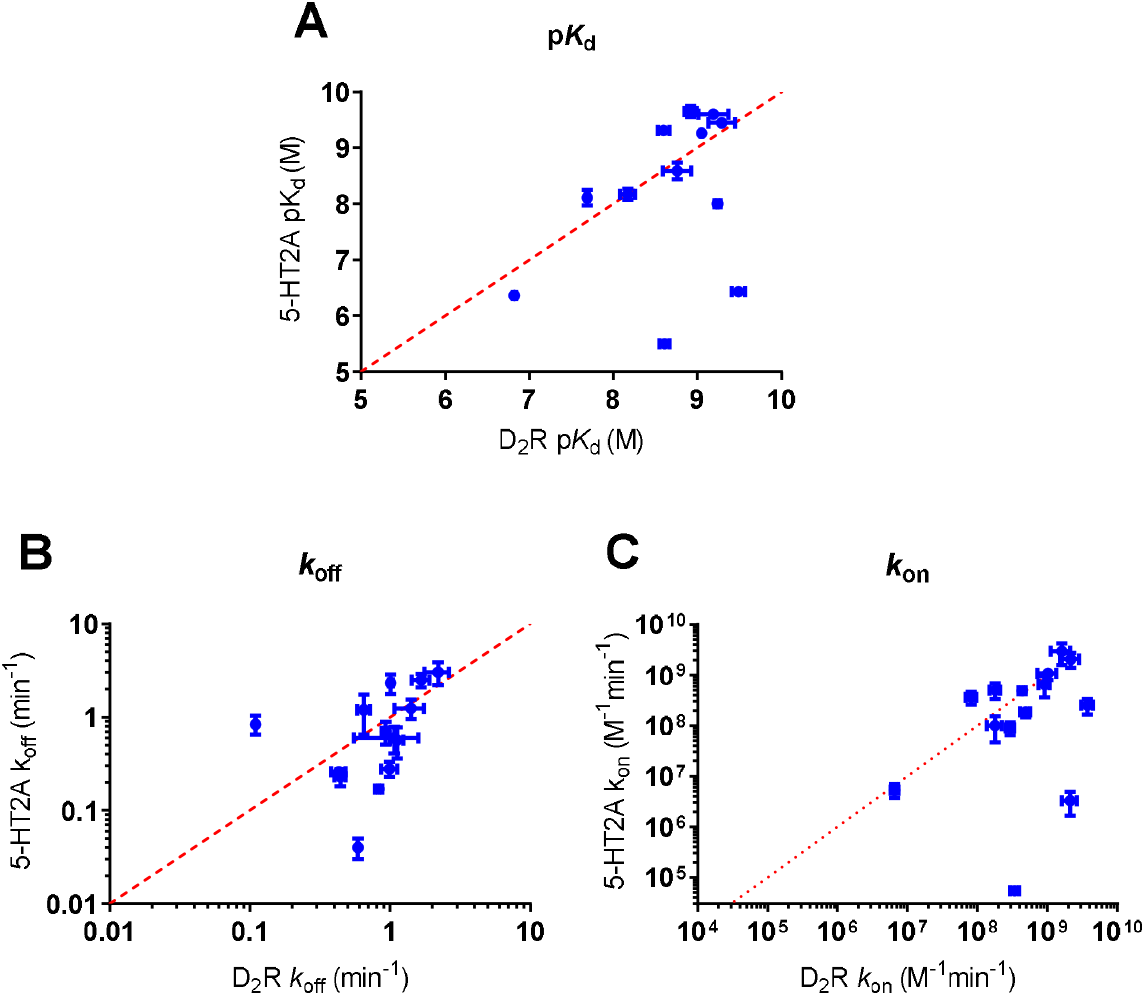
Correlating 5-HT_2A_ and dopamine D_2_R kinetically derived parameters of atypical and typical APDs and reference ligands. Correlation between dopamine D_2_R and 5-HT_2A_R **(A)** p*K*_d_ (-log(*k*_off_/*k*_on_)) values **(B)** *k*_off_ values and **(C)** *k*_on_ values. 5-HT_2A_ p*K*_d_ values were taken from 4F4PP oxalate-red competition association binding experiments as exemplified in Figure 3. All data used in these plots are detailed in Table 1 and supplementary table 1 whilst other D_2_R kinetic and p*K*_d_ values were taken from Sykes et al., 2017. Data are presented as mean ± S.E.M. from at least four experiments.

### Modelling the relative effects of 5-HT2A and dopamine D_2_ receptor rebinding

At least in the brain, target rebinding models are useful predictors of the long-term effects of APDs and reflective of PET studies (Vauquelin and Charlton 2010; Sykes et al., 2017; Seeman 2002). We therefore assessed relative 5-HT_2A_, and D_2_ receptor target occupancy expressed as a function of time by estimating individual drug-receptor target reversal (*k*_r_) at an initial D_2_R striatal occupancy of 80%, based on the rebinding model we reported previously (Sykes et al., 2017). Relative 5-HT_2A_ and D_2_R occupancy (5-HT_2A_: D_2_) was determined by calculating AUC at D_2_R occupancy levels between 80-65% (i.e., those relevant to antipsychotic and EPS through combined serotonergic and D_2_R blockade see Kasper et al., 1999). The relative 5-HT_2A_ and D_2_ receptor target occupancies of a selection of test agents expressed as a function of time are shown in Figure 6A-G with receptor occupancy markers between 80 and 65% marked by the dotted lines. This figure demonstrates the apparent kinetic selectivity of APDs at these two receptors under conditions where rebinding is favoured and its effects on relative receptor occupancy (Vauquelin, 2010; Vauquelin and Charlton, 2010). To achieve a fair comparison, we considered differences in cortical 5-HT_2A_R density relative to striatal D_2_R receptor density. Previous human and animal studies indicate that 5-HT_2A_R numbers are at least 2-3 times greater in the (prefrontal) cortex relative to the striatum (Jafari et al., 2013; Joyce et al., 1991; Muguruza et al., 2013).

**Figure 6.**
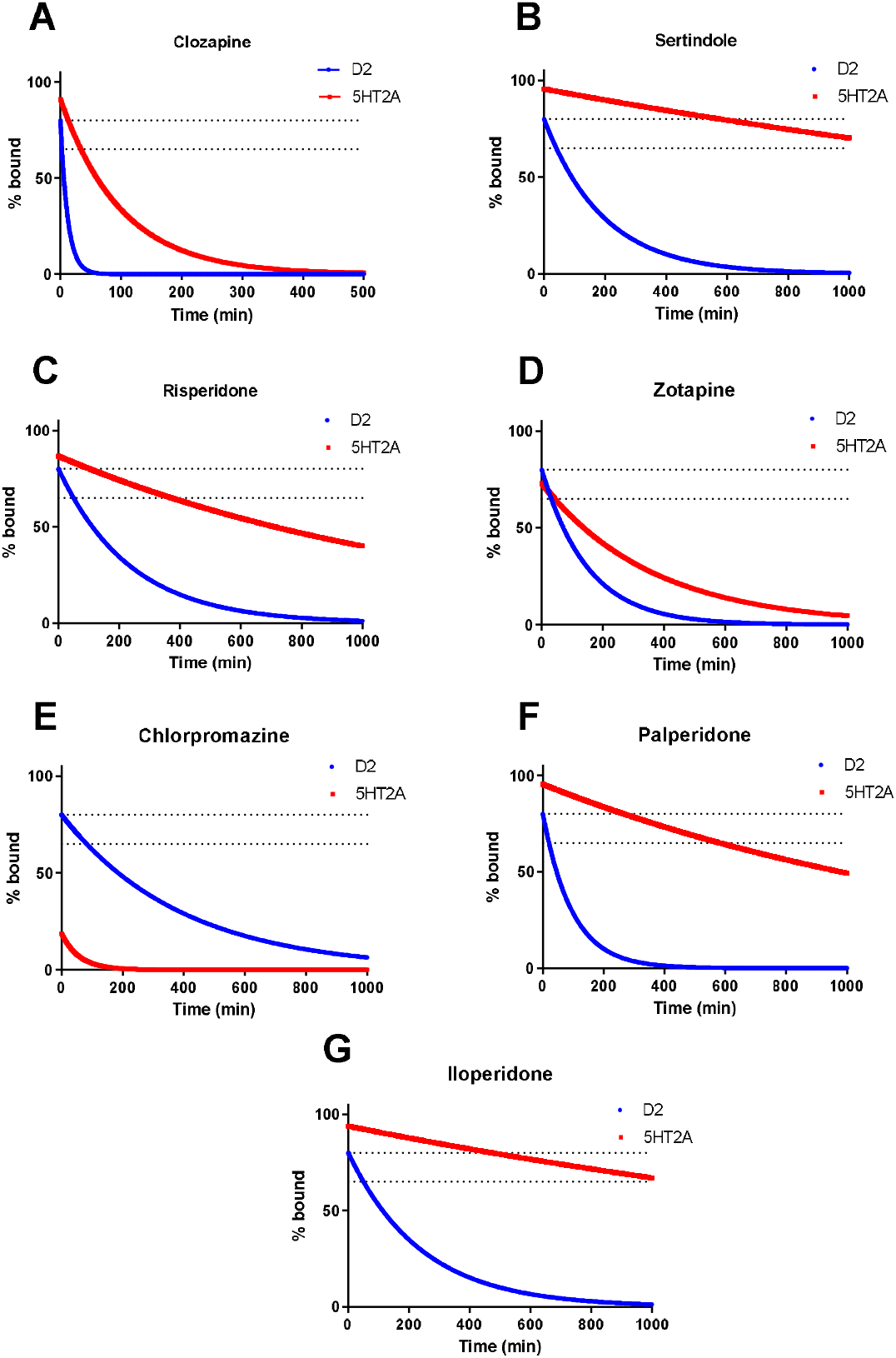
Modelling relative APD D_2_R and 5-HT_2A_ receptor target reversal (*k*_r_) or rebinding. Simulated target reversal rates of clinically relevant APDs from human D_2_ and 5-HT_2A_ receptors, **(A)** clozapine **(B)** sertindole **(C)** risperidone **(D)** zotepine **(E)** chlorpromazine **(F)** paliperidone and **(G)** iloperidone under conditions of limited diffusion based on the association (*k*_on_) and dissociation (*k*_off_) rates determined in competition kinetic binding experiments. All kinetic parameters used to these plots are taken from Table 1 and supplementary table 1 and Sykes et al., 2017 with *k*_r_ values calculated using Equation 4. For simulation purposes the reversal rate *k*_r_ was based on the model of an immunological synapse as detailed in the methods section Modelling drug-receptor re-binding at Dopamine D_2_ and serotonin 5-HT_2A_Rs. The dotted lines represent the levels of dopamine D_2_R occupancy 80-65% thought to be relevant in the occurrence of EPS with clinically used APDs at therapeutic doses (see Kasper et al., 1999).

The original ‘serotonin-dopamine hypothesis’, uses the 5-HT_2A_/D_2_ equilibrium affinity ratio (based on p*K*_i_) to predict the relative incidence of EPS (Meltzer et al., 1989). Our new predictions of individual APD 5-HT_2A_/D_2_ equilibrium affinity ratio (based on 1/*K*_d_) in order of reducing levels of clinically observed EPS are shown in Figure 7A. The 5-HT_2A_: D_2_R AUC ratios for these same compounds in order of reducing levels of clinically observed EPS is shown for comparison in Figure 7B and demonstrates a ceiling effect on the relative 5-HT_2A_: D_2_R receptor occupancy levels suggesting that increases in relative 5-HT_2A_ affinity might not significantly improve any protection provided by a 5-HT_2A_ selective compound. No correlations were observed between clinically observed EPS odds ratio and the 5-HT_2A_: D_2_ affinity and 5-HT_2_: D_2_ *k*_r_ AUC ratios (data not shown).

**Figure 7.**
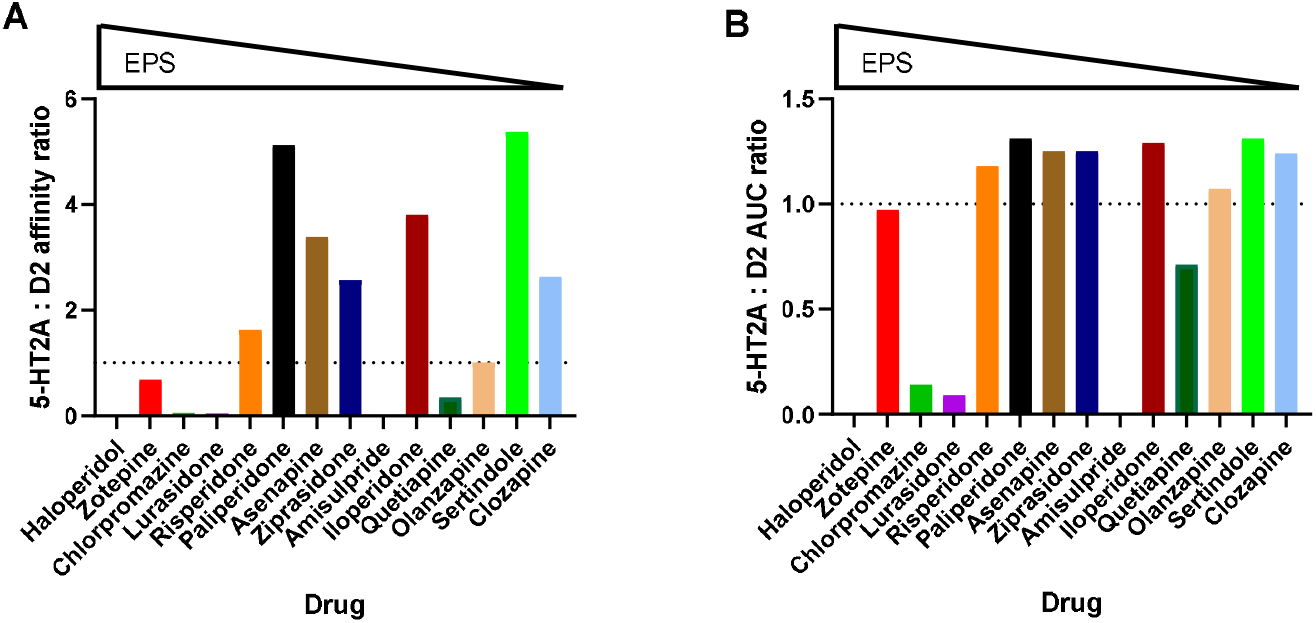
Understanding antipsychotic drug pharmacological parameters and their relationship with clinically observed EPS. **(A)** Antipsychotic drug 5-HT_2A_:D_2_R affinity and **(B)** AUC ratios in order of reducing EPS odds ratio. Relative 5-HT_2A_: D_2_R affinity ratio (based on 1/*K*_d_) and (5-HT_2A_: D_2_R (80-65%) AUC was calculated using measured APD kinetic parameters measured in competition association assays. Relative affinity derived from the kinetic *K*_d_ (*k*_off_/ *k*_on_) and kinetic data used in these plots are detailed in Table 1 and Table 2 and Sykes et al., 2017 with clinical EPS odds ratio data taken from Leucht et al., 2013.

We previously reported a correlation between the rates of compound rebinding and the observed levels of clinically observed EPS. Intriguingly strong dopamine D_2_R re-binders such as haloperidol, zotepine, lurasidone and chlorpromazine, with low 5-HT_2A_: D2 *k*_r_ AUC ratios (i.e <1) showed the widest range in EPS odds ratios potentially since none of these agents (following dosing in patients) is able to positively modulate endogenous release of dopamine via the 5-HT_2A_R to lower overall D_2_R occupancy and reduce the risk of EPS (i.e. no brakes, with EPS effects dependent purely on the level of initial D_2_R blockade, see Figure 8A). Moreover, this plot reveals drugs such as sertindole (and asenapine and ziprasidone) with high 5-HT_2A_: D2 *k*_r_ AUC ratios (i.e >1.0) which show lower levels of EPS in the clinic than expected based on their D_2_R overall rate of binding reversal (see Sykes et al., 2017) (Figure. 8B).

**Figure 8.**
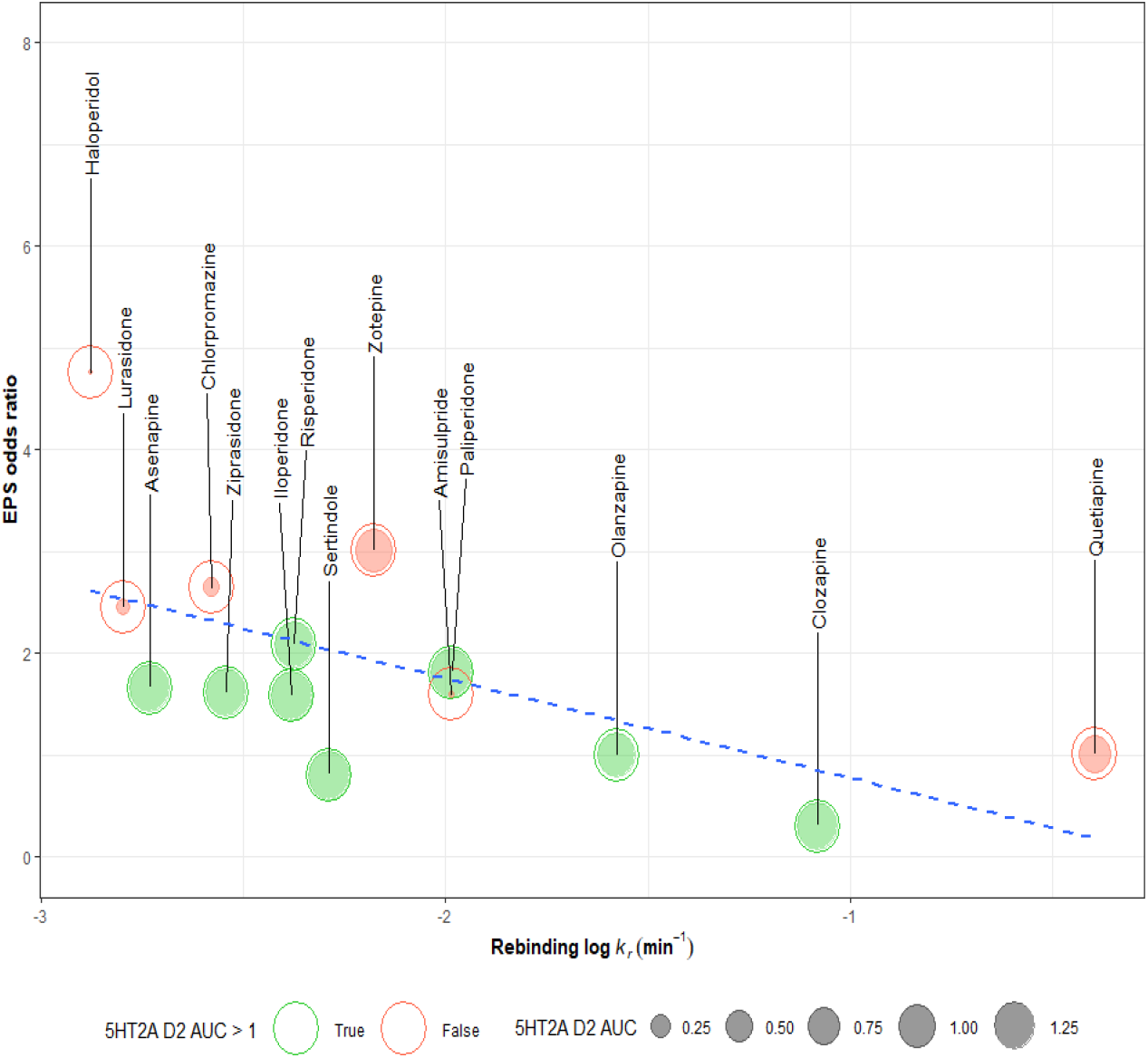
Modelling APD D_2_R rebinding and its relationship with 5-HT_2A_ receptor coverage and consequences for clinically observed ‘EPS. Correlating clinical ‘EPS with the kinetically derived overall reversal rate *k*_r_ at the D_2_R. Correlation plot showing the relationship between log *k*_r_ at D_2_R and EPS odds ratio for all compounds, with 5-HT_2A_R specific compounds highlighted in green. Specificity in this case relates to AUC measurements for target occupancy as determined by relative D_2_R and 5-HT_2A_R target reversal (*k*_r_). All kinetic data used in these plots are detailed in Table 1 and Supplementary Table 1. The relationship between log *k*_r_ and EPS odds ratio is depicted as a trend line, with clinical EPS odds ratio data taken from Leucht et al., 2013.

## Discussion

The novel TR-FRET kinetic assay described herein has enabled us to quantify, for the first time, the kinetic rate constants of a large number of unlabelled clinically used APDs at 5-HT_2A_Rs enabling us to better investigate a role for 5-HT2_2A_R target coverage in the reduced side effect profile of clinically used APDs.

It has been widely reported that the 5-HT_2A_ affinity of certain clinically used atypical antipsychotics is much higher than their corresponding affinities at dopamine D_2_Rs (Horacek et al., 2006; Meltzer et al., 1989; Richelson and Souder, 2000). However, our TR-FRET data revealed a surprisingly small difference in affinity for D_2_ and 5-HT_2A_Rs across the clinically used atypical serotonin-dopamine antagonists (e.g. risperidone, sertindole and iloperidone) and MARTAs studied. Others have reported similar conclusions to our own (Seeman et al., 1997a; Seeman et al., 1997b) although the notion of higher 5-HT_2A_ affinity for atypical APDs persists (Langlois et al., 2012; Meltzer and Massey, 2011). Potential reasons for these discrepancies include depletion of high-affinity radioligands such as [^3^H]-spiperone potentially leading to underestimated D_2_R tracer and therefore unlabelled compound affinities, in conjunction with differences in assay temperature and buffer salt concentration (Seeman et al., 1997a).

It is suggested that typical APDs block dopamine stimulated output via their inhibitory action at D_2_Rs throughout the brain. In contrast, atypical antipsychotics with 5-HT_2A_ inhibitory effects have more complex net actions on dopamine activity and are able to not only block dopamine activity at D_2_Rs but also potentially increase dopamine release via 5-HT_2A_R blockade and thus promote dopamine signalling in certain select regions of the brain (Horacek et al., 2006; Lucas and Spampinato, 2000; Stahl, 2008). Consequently, it has been suggested that these agents will have therapeutic actions not just on the positive symptoms of schizophrenia but also on negative, cognitive and affective symptoms (Horacek et al., 2006). Also, a significant reduction in the incidence of EPS might be expected through increased dopamine release in the striatum (Horacek et al., 2006) and potentially a reduction in hyperprolactinemia through increased dopamine release at the level of the hypothalamus (Compton and Miller, 2002).

Target occupancy in the brain is not well described by traditional models of receptor binding which assume free diffusion. Thus, to compare relative dopamine D_2_R and 5-HT_2A_R occupancy we have employed a model which accounts for rebinding of molecules in conditions of limited diffusion (Sykes et al., 2017; Vauquelin and Charlton, 2010). The relative 5-HT_2A_ and D_2_ receptor target occupancies of selected test agents under the influence of this model and expressed as a function of time is shown in Figure 6A-E. What is apparent is the wide variation in relative occupancy of atypical APDs at these two receptors when plotted as a function of time. This figure illustrates the potential of rebinding to dictate both drug-receptor selectivity and control relative receptor occupancy under conditions of limited diffusion, with certain molecules such zotepine, which historically has been classed as both atypical and typical showing less occupancy at the 5-HT_2A_R whereas other agents such as clozapine appear to be more balanced and relatively short-acting at both receptors. In contrast, there are atypical compounds that show a high level of (long-lasting) occupancy at both receptors such as sertindole, a compound known for its potent and long-lasting effects at 5-HT_2A_R (Hyttel et al., 1992; Sánchez et al., 1991).

We compared our new drug 5-HT_2A_R kinetic and binding data with clinical data that quantified the level of EPS with a diverse group of clinical APDs. The clinical data were taken from a relatively comprehensive multiple-treatment meta-analyses of antipsychotic drug efficacies (Leucht et al., 2013). Whilst relative affinity for the 5HT_2A_: D_2_ receptors appears to modulate absolute EPS the suggestion that higher 5-HT_2A_ affinity alone of atypical APDs (such as clozapine) determines their reduced side EPS profile is not supported by our kinetic (or equilibrium) affinity data, see Figure 7A. 5-HT_2A_: D_2_ receptor affinity ratio estimates provide a snapshot of receptor occupancy at equilibrium, whereas AUC measurements provide a relative measure of target occupancy over time and consequently should better predict their action in the body (see Figure 7B). As noted before the clear outlier in this scheme is the lower affinity D_2_R antagonist amisulpride which has negligible affinity for the 5-HT_2A_R (Kapur and Seeman, 2001).

This analysis of 5-HT_2A_R binding kinetics has effectively identified two distinct groups of APDs which we represent graphically in Figure 8. One group is APDs whose clinical EPS effects are influenced mainly by their effective D_2_R reversal rate (*k*_r_), and another group whose EPS are further modulated by higher levels of 5-HT_2A_R occupancy. These results strongly suggest that the relative level of dopamine output dictated by 5-HT_2A_: D_2_R occupancy over time could have an impact on observed EPS and may partly explain the imperfect correlation we observed with the binding reversal rate (*k*_r_) in our previous study (Sykes et al., 2017). Interestingly sertindole’s effects appear to be underestimated by this model potentially because of its own uniquely slow off-rate from the 5-HT_2A_R.

The ability of this compound to produce insurmountable antagonism is highlighted in Figure 4B. In direct contrast to sertindole, clozapine’s effects are largely reversible (minimal reduction in E_max_) under identical conditions (Figure 4A). Hypothetically augmented dopamine release due to sustained 5-HT_2A_ residency could partially compensate for high striatal D_2_R occupancy as a result of robust receptor rebinding and a relatively slow dissociation rate (*k*_off_).

Atypicality in terms of on target EPS seems to be attributable to two independent pharmacological properties of APDs; rapid reversal of D_2_ receptor occupancy in the striatum due to reduced APD rebinding potential (as is the case for compounds with low 5-HT_2A_: D_2_R coverage such amisulpride and olanzapine) and high 5-HT_2A_: D_2_R coverage the extreme example being sertindole.

Our discovery that lurasidone has a relatively slower dissociation rate (*k*_off_) from the D_2_R seems to further confirm the key role of association rate (*k*_on_) over dissociation rate in the prevalence of APD EPS with agents such as haloperidol exhibiting much higher levels of clinically observed EPS despite its much faster dissociation rate.

These results suggest that the balance between 5-HT_2A_R occupancy and D_2_R occupancy is the key to the reduced side effect profile of sertindole with increased dopamine release potentially compensating for enhanced striatal D_2_R occupancy as a result of increased receptor rebinding. The dampening effects of sustained 5-HT_2A_R occupancy would appear limited, due to the nonlinear relationship between affinity and AUC ratios which demonstrates a ceiling effect (see Supplementary Figure 2).

It is of interest to this discussion that the in vivo dopamine D_2_R occupancy profile of the atypical antipsychotic risperidone appears relatively flat in comparison to haloperidol, which is dopamine D_2_R selective (Kusumi et al., 2015; Natesan et al., 2006). This could be an indication that the high 5-HT_2A_R occupancy of risperidone can modulate D_2_R occupancy, potentially as a result of increased dopamine release in the striatum. Indirect evidence for this scenario comes from a PK-PD model produced through studying the effects of atypical and typical APDs in patients during a 6-week clinical trial (Pilla Reddy et al., 2012). This study by Reddy and colleagues, confirms that the probability of experiencing EPS, as D_2_R receptor occupancy increases, is higher for the D_2_R selective typical APDs such as haloperidol compared to atypical antipsychotics with significant 5-HT_2A_R affinity such as ziprasidone. This model and these findings are consistent with a previous study of antipsychotic-induced extrapyramidal symptoms which utilised a D_2_R receptor occupancy model incorporating endogenous dopamine release, the result of 5-HT_2A_ inhibition (Matsui-Sakata et al., 2005).

One alternative theory suggests that 5-HT_2A_R blockade could reduce the release of dopamine in the limbic system and therefore the level of dopamine receptor blockade required in this area of the brain is likely to also be reduced (Howell and Cunningham, 2015). This would fit with a reduction in the incidence of positive symptoms by 5-HT_2A_R selective ligands and could also partly explain why clozapine is so effective in uncontrolled schizophrenia. As a knock-on effect, a reduction in the absolute level (%) of receptor blockade observed in the striatum, due in part to limbic selectivity, might be expected. The outcome, if such a mechanism is correct, should be a reduction in the likelihood of patients experiencing EPS, effectively reinforcing the key role of the 5-HT_2A_R in APD atypicality.

It must be acknowledged that our analysis does not suggest an exclusive role for the 5-HT_2A_R in the reduced EPS effects of the atypical APD sertindole or the other serotonin-dopamine antagonists. Both sertindole and asenapine have been shown to have almost identical affinity for the serotonin 5-HT_2C_ receptor (Schotte et al., 1996; Shahid et al., 2009), and so it is entirely possible that this often overlooked receptor is more important in dictating the absolute levels of drug-induced EPS (Di Giovanni and De Deurwaerdere, 2016; Gardell et al., 2007; Reavill et al., 1999). Interestingly in this regard, APDs with a lower affinity for the 5-HT_2C_ receptor relative to the 5-HT_2A_R would appear to show higher levels of EPS e.g., Iloperidone, risperidone, paliperidone and ziprasidone (Herrick-Davis et al., 2000; Kalkman et al., 2001; Mauri et al., 2017). Therefore, the role of the 5-HT_2C_ receptor in reducing EPS remains an open question and one which is worthy of future study.

In conclusion, higher affinity atypical drugs with a significant affinity for the 5-HT_2A_Rs such as asenapine and sertindole represent an advance in reducing the on-target EPS profile of APDs relative to conventional APDs. The lack of EPS observed with sertindole suggest slow dissociation from serotonergic receptors could be advantageous in terms of this profile. However, none of these agents is as effective as clozapine both in terms of clinical efficacy and reducing EPS.

## Abbreviations

CHO: Chinese hamster ovary
HBSS: Hanks’ balanced salt solution
[NSB: non-specific binding

## Acknowledgments

This work was carried out at the University of Nottingham. We acknowledge the contribution of Dr Nicholas Holiday and Dr Lisa Stott who contributed to the generation of plasmids and cell lines used to create the 5-HT2A and D_2_ receptor binding assays.

## Supplementary Figures

**Supplementary Figure 1.**
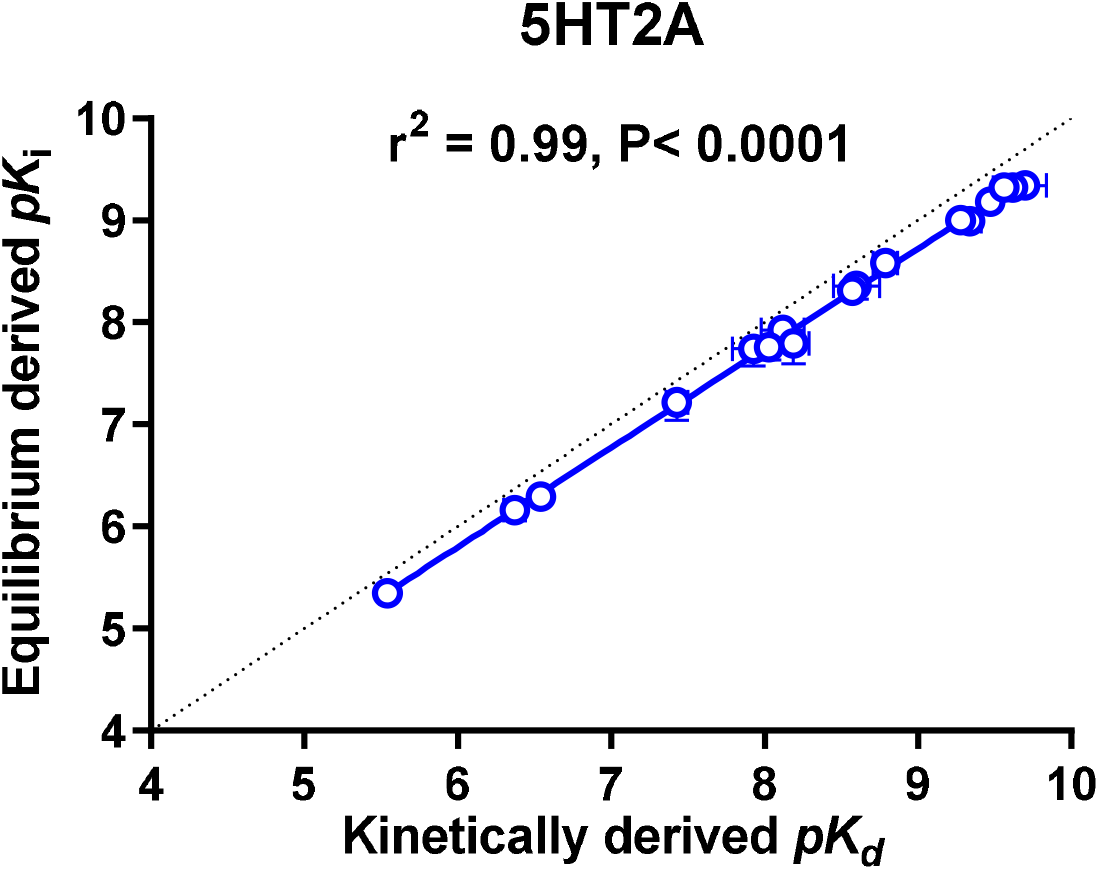
Correlating serotonergic receptor equilibrium and kinetically derived parameters of atypical and typical APDs and reference ligands. Correlation between kinetically derived p*K*_d_ (-log(*k*_off_/*k*_on_)) and 5-HT_2A_ p*K*_i_ values for the test ligands. p*K*_i_ values were taken from competition binding experiments at equilibrium. All data used in these plots are detailed in Table 1. Data are presented as mean ± S.E.M. from 4-5 experiments. The relationship between p*K*_i_ and kinetically derived p*K*_d_ was assessed using a two-tailed Pearson’s correlation coefficient (r^2^). A *P* value of 0.05 was used as the cut off for statistical significance.

**Supplementary Figure 2.**
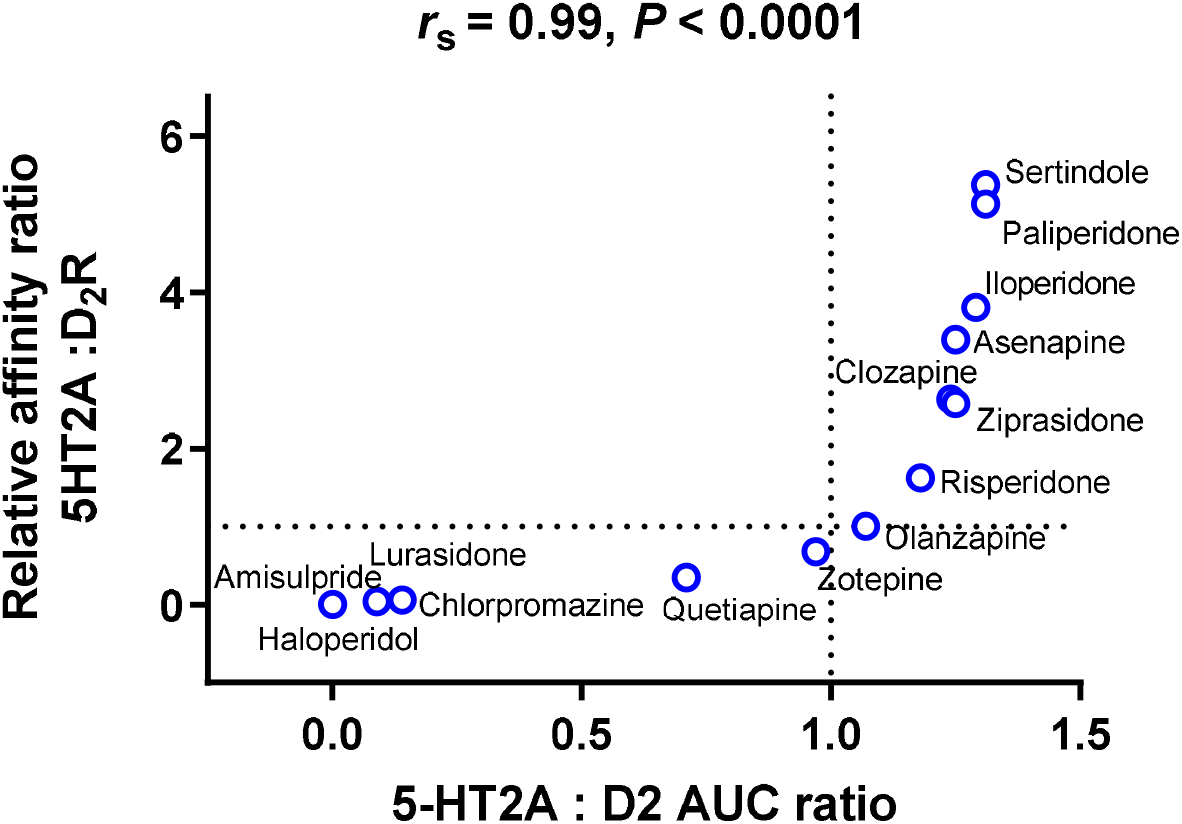
Relationship between 5HT_2A_: D_2_R AUC ratios and 5HT_2A_: D_2_R AUC affinity ratios. Relative affinity derived from the kinetic *K*_d_ (*k*_off_/ *k*_on_) and kinetic data used in these plots are detailed in Table 1 and Table 2 and Sykes et al., 2017 with clinical data taken from Leucht et al., 2013. Kinetic data are presented as mean ± S.E.M. from at least four experiments and clinical data as the odds ratio for EPS with associated upper credible interval where indicated. The relationship between two variables was assessed using a two-tailed Spearman’s rank correlation allowing the calculation of the correlation coefficient, r_s_. A P value of 0.05 was used as the cut off for statistical significance.

**Supplementary table 1.** Kinetic binding parameters of unlabelled clinically used APDs and reference antagonists for human D_2L_ receptors and their historical classification as atypical APDs.

|  | *k<sub>off</sub>* (*min*<sup>-1</sup>) | *k<sub>on</sub>* (*M*<sup>-1</sup>*min*<sup>-1</sup>) | *Dissociation*<br>*t*<sub>1/2</sub> (*min*) | *pK<sub>d</sub>* | *pK<sub>i</sub>* | *n* | Receptor occupancy<br>reversal rate or *k<sub>r</sub>*<br>(*min*<sup>-1</sup>) |
| --- | --- | --- | --- | --- | --- | --- | --- |
| **SGA/Atypical** |  |  |  |  |  |  |  |
| Lurasidone | 0.11 ± 0.00 | 2.95 ± 0.40 x10<sup>8</sup> | 6.36 | 9.41 ± 0.09 | 9.57 ± 0.14 | 4 | 0.005405 |
| lloperdone | 0.99 ± 0.13 | 1.03 ± 0.30 x10<sup>9</sup> | 0.79 | 8.96 ± 0.10 | 8.99 ± 0.10 | 6 | 0.03128 |

Data are mean ± S.E.M. for the number of observations (n) stated, each performed in singlet.

